# TCF4 induces enzalutamide resistance via neuroendocrine differentiation in prostate cancer

**DOI:** 10.1101/560821

**Authors:** Geun Taek Lee, Won Tae Kim, Young Suk Kwon, Ganesh Palapattu, Rohit Mehra, Wun-Jae Kim, Isaac Yi Kim

## Abstract

In treating patients with castration resistant prostate cancer (CRPC), enzalutamide, the second-generation androgen receptor (AR) antagonist, is an accepted standard of care. However, clinical benefits are limited to a median time of 4.8 months because resistance inevitably emerges. To determine the mechanism of treatment resistance, we carried out a RNA sequence analysis and found increased expression levels of neuroendocrine markers in the enzalutamide-resistant LNCaP human prostate cancer (CaP) cell line when compared to the parental cell line. Subsequent studies demonstrated that TCF4, a transcription factor implicated in Wnt signaling, mediated neuroendocrine differentiation (NED) in response to enzalutamide treatment and was elevated in the enzalutamide-resistant LNCaP. In addition, we observed that PTHrP mediated enzalutamide resistance in tissue culture and inducible TCF4 overexpression resulted in enzalutamide-resistance in a mouse xenograft model. Finally, small molecule inhibitors of TCF4 or PTHrP partially reversed enzalutamide resistance in CaP cells. When tissues obtained from men who died of metastatic CaP were examined, a positive correlation was found between the expression levels of TCF4 and PTHrP. Taken together, the current results indicate that TCF4 induces enzalutamide resistance via NED in CaP.

## Introduction

Prostate cancer (CaP) is the most common non-cutaneous cancer diagnosed among men and the second leading cause of male cancer deaths in the United States (1). In 2017, it is estimated that 26,730 men died from CaP. Although radiation and surgery are quite effective for localized disease, approximately 30% eventually recur following a definitive therapy. More importantly, there is no effective cure for men who present with metastatic CaP as the 5-year relative survival rate is only 29% (1). In patients with a metastatic disease, medical or surgical castration is generally the accepted first-line therapy. Yet, castration-resistant prostate cancer (CRPC) eventually emerges with a median time of 18-24 months (2, 3). Once CRPC develops, secondary hormonal manipulation, immunotherapy, and chemotherapy are marginally effective and the average life expectancy is ~5 years (4, 5).

Enzalutamide is a FDA-approved second-generation androgen receptor (AR) antagonist that blocks ligand binding, nuclear translocation, DNA binding, and coactivator recruitment of ARs (6). In multiple clinical trials, enzalutamide has been shown to prolong overall and progression-free survival, improve patient-reported quality of life, and delay the development of skeletal-related complications in men with metastatic CRPC who are chemotherapy naïve or have previously received docetaxel (7-9). However, despite the significant initial therapeutic benefits of enzalutamide, resistance inevitably occurs with a median time of 4.8 months. Although the precise mechanistic details underlying the emergence of enzalutamide resistance is largely unknown, the activation of adaptive survival pathways in an androgen-depleted environment is likely important.

TCF4, also known as transcription factor 7-like 2 (TCF7L2), is an important effector of the canonical Wnt signaling pathway (10). Although dysregulated Wnt signaling has extensively been linked to CaP cells (11-13), TCF4 has not been demonstrated directly to play a role in CaP progression. In an effort to identify critical pathways involved in enzalutamide resistance, we developed multiple enzalutamide-resistant human CaP cell lines and found that TCF4 mediates enzalutamide resistance in CaP cells by inducing neuroendocrine differentiation (NED) via a Wnt-independent mechanism.

## Materials and Methods

### Cell culture

Human prostate cancer cell lines, LNCaP, 22Rv1, and VCaP were obtained from the American Type Culture Collection (Manassas, VA) and maintained in the standard culture media: RPMI-1640 supplemented with 10% fetal bovine serum (FBS). TCF4 cDNA (Origene, Cat# 224345, Rockville, MD) was cloned into pLenti4/V5-Dest expression vector (ThermoFisher, Waltham, MA). After making pLenti4/V5-Dest/TCF4 viral supernatant with the Optimized Packaging Mix, the supernatant was added to the Virapower T-Rex LNCaP. After selection with Zeocin and Blasticidin, TCF4-inducible LNCaP (TCF4/LNCaP) were selected and the expression of TCF4 was screened using quantitative PCR (QPCR). Selected clones were cultured in RPMI-1640/10% FBS media containing 100 mg/ml of Blasticidin and Zeocin (Life Technologies, Carlsbad, CA, USA). To establish enzalutamide resistant CaP cell lines, cells were treated continuously with 10μM enzalutamide (Selleckchem, Houston, TX, USA, Cat# S1250). After three months, stable cell lines were established (LNCaP-EnzR, VCaP-EnzR, and 22Rv1-EnzR). Unless specified, the standard culture media for these three enzalutamide-resistant cell lines included 10 μM enzalutamide. The β-catenin degradation activator was purchased from (Selleckchem, Houston, TX, USA, Cat# S1180).

### Mice study

All mice and experimental procedures were conducted using protocols approved by and in accordance with the Rutgers Cancer Institute of New Jersey Institutional Animal Care and Use Committee approval (PROTO999900168) and the National Institutes of Health Guide for the Care and Use of Laboratory Animals. For anesthesia, 3% isoflurane gas inhalation method was used. Rag2-/-, γc-/- mice were purchased from The Jackson Laboratory (Bar Harbor, Maine, USA). The study was approved by the Institutional Animal Care and Use Committee at the Rutgers University (New Brunswick, NJ). Where indicated, surgical castration was carried out via bilateral orchiectomy. Tumor size was measured using calipers and tumor volume was calculated using the formula: tumor volume= length x width^2^ /0.361. Doxycycline-inducible TCF4-expressing cells were subcutaneously injected into five mice per group. Mice were supplied with water containing 2 mg/ml of doxycycline (Sigma-Aldrich, Saint Louis, MI, USA) in 5 % sucrose to induce TCF4.

To explore the therapeutic implications of targeting TCF4/NED pathway, LNCaP-EnzR was injected into forty Rag2-/-, γ_c_-/- immunodeficient mice. When tumors reached an average size of 3 mm in diameter, all animals were surgically castrated and divided into four groups of ten mice each. TCF4/β-catenin interaction was disrupted with PKF118-310 while PTHrP was blocked with PTHrP_(7-34)_. PKF118-310 was purchased from Millipore (Burlington, MA, Cat# 219331) and PTHrP_(7-34)_ Bachem (Torrance, CA, Cat# H-9100.0500). Mice were injected daily with the PKF118-310 (0.85 mg/kg, intraperitoneal) and/or PTHrP_(7-34)_ (0.2 mg/kg, subcutaneous) for four weeks. Where indicated, enzalutamide was administered daily via oral gavage at 10 mg/kg in 1% carboxymethyl cellulose, 0.1% Tween-80, and 5% DMSO (14-16). Tumor volume and body weight were measured weekly. Four weeks after castration, all animals were sacrificed and tumors were harvested and analyzed for the expression of TCF4 and PTHrP using immunofluorescence microscopy. Animals were euthanized by CO_2_ asphyxiation after PKC118-310 and/or PTHrP_(7-34)_ treatment were completed.

### RNA sequencing

RNA was purified using DirectZol RNA purification kit (Zymoresearch, Irvine, CA, Cat# R2060) from LNCaP, charcoal stripped FBS (cFBS)-resistant LNCaP (LNCaP-cFBSR), and LNCaP-EnzR. Then, RNA sequencing was performed by Macrogen, Inc. (Washington D.C., USA) using purified RNA.

### Transient Transfections

One μg of a plasmid containing TCF4 cDNA (Origene, Cat# 2243345, Rockville, MD) was transfected into indicated prostate cancer cell lines on a 6-well plate. Three μl of lipofectamine 3000 (ThermoFisher, Waltham, MA, USA, Cat# L3000015) was used for each transfection.

### Quantitative RT-PCR and PCR

Total RNA was isolated with TRIzol LS reagent (Thermo Fisher Scientific, Waltham, MA, USA), and 1-2 μg of total RNA was used for synthesizing cDNA with High-Capacity cDNA Reverse Transcription Kit (Thermo Fisher Scientific, Waltham, MA, USA). cDNA was then used for Q-PCR in a StepOnePlusTM (Applied Biosystems, Foster City, CA, USA) with SYBR Green ROX qPCR Mastermix (QIAGEN, Valencia, CA, USA). Sequences of PCR primers used are as follows: human QPCR ChgA (forward: GAAGAGGAGGAGGAGGAGGA, reverse: CACTCAGGCCCTTCTCTCTG); human QPCR β-actin (forward: AGAGCTACGAGCTGCCTGAC, reverse: AGCACTGTGTTGGCGTACAG); human QPCR TCF4 (forward: CCTGGCTATGCAGGAATGTT, reverse: CAGGAGGCGTACAGGAAGAG); human QPCR NSE (forward: GGTCCAAGTTCACAGCCAAT, reverse: CAGTTGCAGGCCTTTTCTTC); human QPCR PTHrP (forward: CAAGATTTACGGCGACGATT, reverse: GAGAGGGCTTGGAGTTAGGG)

### Immunoblot analysis

CaP cells were collected and lysed with lysis buffer (20 mM Tris-HCl (pH 7.5), 150 mM NaCl, 1 mM Na_2_EDTA, 1 mM EGTA, 1% Triton, 2.5 mM sodium pyrophosphate, 1 mM beta-glycerophosphate, 1 mM Na_3_VO_4_, 1 µg/ml leupeptin) containing 1 mM phenylmethylsulfonyl fluoride (PMSF). Cell lysates were centrifuged and protein in the supernatant was measured. After separating of 25-50μg protein using SDS-PAGE, samples were incubated with TCF4 (Cell Signaling Technologies, Danvers, MA, USA), ChgA (Abcam, Cambridge, MA, USA), NSE (Abcam, Cambridge, MA, USA), PTHrP (Abcam, Cambridge, MA, USA), POU2F2 (Sigma-Aldrich, Allentown, PA, USA) or β-actin (Sigma-Aldrich, St. Louis, MO, USA) antibodies. For ChgA, NSE, PTHrP, and POU2F2 immunoblots, 1:1000 diluted antibody solutions in 5% skim milk was used. For the β-actin immunoblot, 1:10000 diluted antibody solution was used. All membranes were incubated overnight at 4°C. Following the incubation with appropriate secondary antibody, immunoblot was analyzed using SuperSignal West Femto Maximum Sensitivity Substrate (ThermoFisher Scientific, Waltham, MA, USA).

### TCF4 knockdown

TCF4 MISSION shRNA was purchased from Sigma-Aldrich (St. Louis, MO, USA, Cat# SHCLNG-NM_003199). TCF4 shRNA lentiviral supernatant was generated with ViraPower Lentiviral Packaging Mix (Thermo Fisher Scientific, Waltham, MA, USA) and used to infect LNCaP cells. The expression of TCF4 was analyzed using QPCR and immunoblot as described above.

### Analysis of human TMAs and murine tumors

CRPC tissue microarray (TMA) was obtained from the University of Michigan’s rapid autopsy program. The rapid autopsy program was supported by Specialized Program of Research Excellence in Prostate Cancer (SPORE, National Cancer Institute grant CA69568) and the Prostate Cancer Foundation (17). The University of Michigan Prostate Rapid Autopsy Program protocol meets all the institutional requirements as indicated by the support from the SPORE grant and the protocol is not subject to IRB-MED approval. The array contains 51 CRPC samples as well as 16 benign prostate tissues and 12 localized prostate cancer tissues for controls. TMA and mouse tumor slides were scanned using the Olympus VS120 Florescence/Bright-field whole slide scanner (Olympus Scientific Solutions Americas Corp., Waltham, MA, USA) after staining with the appropriate antibody. All scanned cores or slides were individually quantified using NIH ImageJ V1.50i (NIH, Bethesda, MD, USA). Values were represented as mean fluorescence intensity (MFI). Antibodies to TCF4 (Millipore) and PTHrP (Abcam, Cambridge, MA) were obtained from commercial sources.

### Statistical analysis

Statistical significance was calculated using the Student’s t-test for paired comparisons of experimental groups and, where appropriate, by Wilcoxon rank sum test, and by 2-way ANOVA. *In vitro* experiments were repeated a minimum of three times. Continuous variables were expressed as mean ± standard deviation (SD).

## RESULTS

### Enzalutamide-resistant human CaP cell line exhibits NED

To investigate the mechanism of enzalutamide resistance in CaP cells, an enzalutamide-resistant human CaP cell line was initially generated by continuously treating LNCaP with 10 μM enzalutamide in RPMI-1640 supplemented with 10% charcoal stripped fetal bovine serum (cFBS). After three months, cells began to proliferate consistently and were designated as LNCaP-EnzR. Simultaneously, LNCaP cultured chronically under cFBS was also generated (LNCaP-cFBS). With these cells, RNAseq followed by an unsupervised data analysis was carried out. When compared to the LNCaP-cFBS, there was no obvious differences in AR signaling related gene expression levels. However, NED markers such as chromogranin A (CHGA, ChgA), neuron-specific enolase (ENO2, NSE), and PTHrP (PTHLP) were significantly higher in LNCaP-EnzR (Fig 1A). A similar pattern of increased NED was observed in LNCaP-cFBS when compared to the parental line maintained in the standard media (RPMI1640/10% FBS) (Fig 1B). To validate these observations, we treated the human prostate cancer cell lines LNCaP and 22Rv1 with increasing concentrations of enzalutamide (0-10 μM) under an androgen-deprived condition (RPMI-1640/10% cFBS) for 48 hours. The results demonstrated that enzalutamide increased expression levels of ChgA, NSE, and PTHrP protein (Fig 1C) and mRNA (Fig 1D) in a concentration-dependent manner.

**Figure 1.**
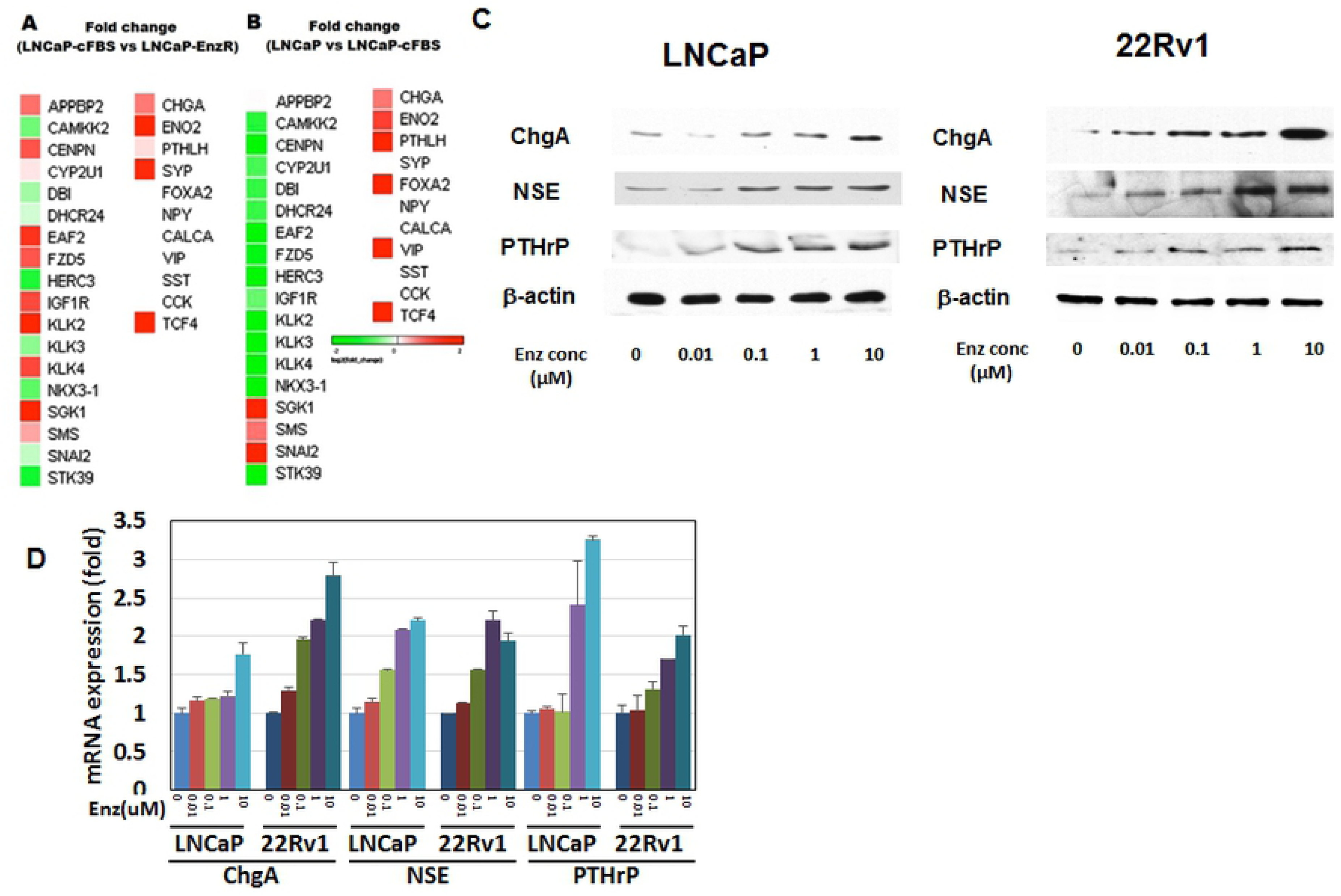
TCF4 mediates NED in human prostate cancer cell lines. **A.** Unsupervised comparison of transcriptome of LNCaP cultured in FBS vs charcoal-stripped FBS (cFBS) was carried out. **B.** Comparison of transcriptome between LNCaP-cFBS and LNCaP-Enz (resistant to enzalutamide). Markers of NED (ChgA, NSE, and PTHrP) were significantly increased in LNCaP-EnzR. **C**. Human prostate cancer cell lines LNCaP and 22Rv1 were treated with increasing concentrations of enzalutamide (0-10 μM) under an androgen-deprived condition (RPMI-1640/10% cFBS) for 48 hours. Immunoblot demonstrated that NED markers (ChgA, NSE, and PTHrP) were induced by enzalutamide. **D.** QPCR demonstrated that NED marker (ChgA, NSE, and PTHrP) mRNA expression levels increased also after treatment with enzalutamide at the indicated concentrations (0-10 μM) for 48 hours.

### TCF4 mediates NED and enzalutamide resistance in human CaP cell lines

Based on the observation that the mRNA levels of NED markers are induced by enzalutamide and increased in LNCaP-EnzR, we hypothesized that the anti-androgen may regulate a common transcription factor that regulates NED. To test this concept, we carried out a bioinformatics-based analysis of the ChgA, NSE, and PTHrP promoters to identify common transcription factors that potentially bind to all three NED markers (alggen, http://alggen.lsi.upc.es/). This effort identified consensus sequences for binding of five transcription factors within the promoters of NED markers: TCF4, POU2F2 (Oct2.1), MRF-2, LCR-F1, and MBF-1 (EDF-1). Quantitative PCR (qPCR) confirmed that mRNA levels of TCF4 and POU2F2 were significantly higher in LNCaP-EnzR when compared to the parental cell line (data not shown). This QPCR result was consistent with our transcriptome analysis that showed increased TCF4 expression levels in LNCaP-EnzR (Fig 1A). To assess the role of TCF4 and POU2F2 on NED, three androgen-responsive human CaP cell lines (LNCaP, 22Rv1, and VCaP) were transiently transfected with TCF4 and POU2F2. Only cells overexpressing TCF4 demonstrated significant increase in the mRNA levels of the NED markers ChgA, NSE, and PTHrP (Fig 2A). In addition, it was found that TCF4-expressing cells were more resistant to enzalutamide 10 μM treatment as the cell number was nearly double that of the control after three days of culture (Fig 2B). This resistance was not merely due to differentiation as TCF4-expressing cells continued to proliferate in the presence of enzalutamide. When the two transcription factors were knocked down in LNCaP and 22Rv1 with shRNA, enzalutamide no longer induced the expression of ChgA, NSE, and PTHrP proteins only when TCF4 expression was blocked (Fig 2C). The kinetics of enzalutamide-induced TCF4 and NED markers expression revealed that the increase in TCF4 mRNA preceded that of the NED markers by approximately eight hours in tissue culture (Fig 2D). These results collectively demonstrate that TCF4 mediates the expression of NED markers in human CaP cell lines.

**Figure 2.**
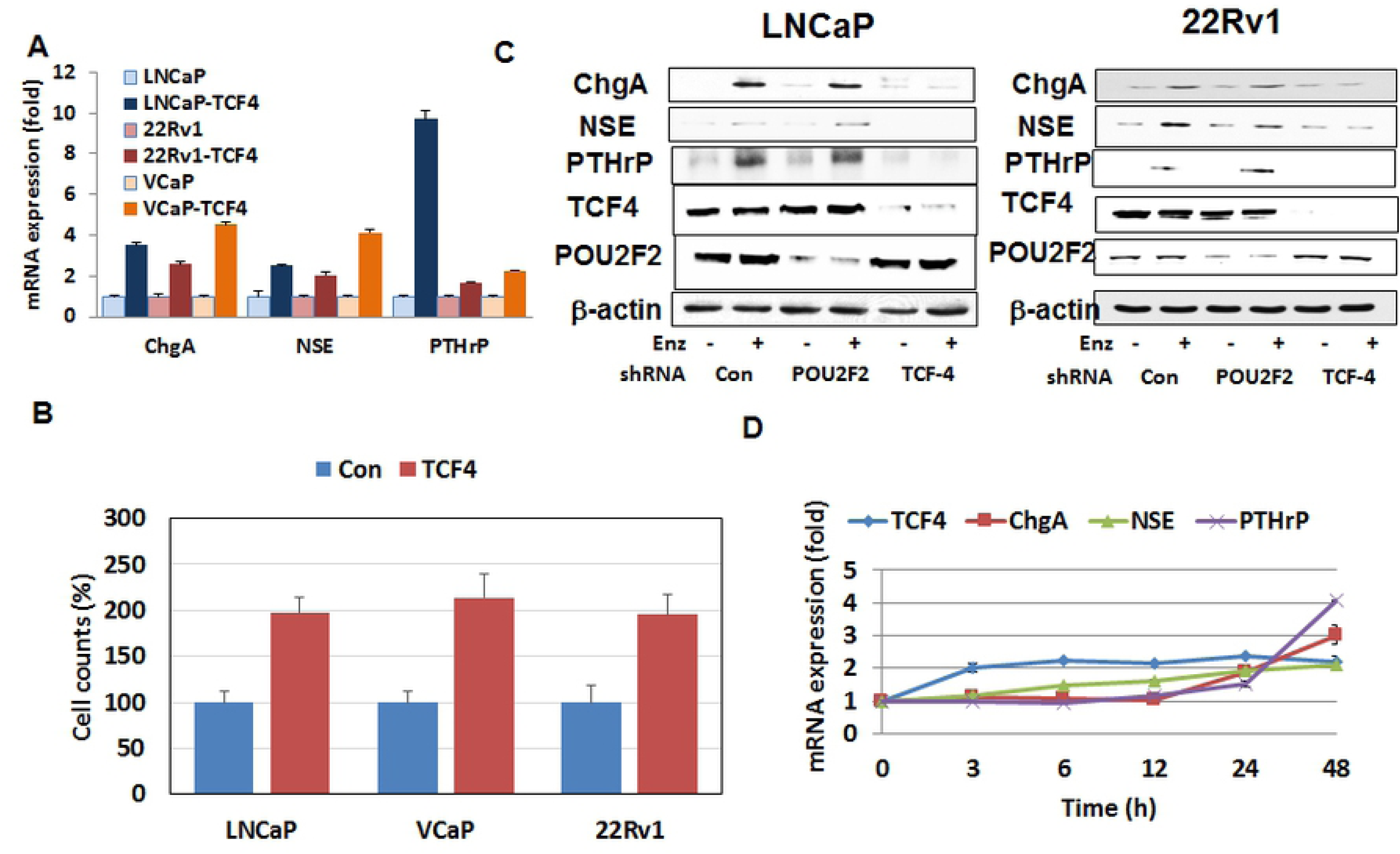
TCF4 mediated enzalutamide resistance in human prostate cancer cell lines. **A.** TCF4 cDNA was transiently transfected into LNCaP, 22Rv1, and VCaP using lipofectamine. Cells were analyzed 48 hours after transfection. The results demonstrated that the overexpression of TCF4 induced the mRNA expression levels of LNCaP, 22Rv1, and VCaP. As control, parental lines transfected with the plasmid backbone was used. **B**. In addition to increasing NED, overexpression of TCF4 increased the cellular proliferation rate of LNCaP, VCaP, and 22Rv1. The result shows cell counts at 72 hours after transfection. **C.** LNCaP and 22Rv1 were treated with enzalutamide for 48 hours. Where indicated, TCF4 or the control POU2F2 expression was silenced using shRNA approach. Increased protein levels of neuroendocrine markers following enzalutamide (enz) treatment was blocked by TCF4 shRNA in LNCaP and 22Rv1. POU2F2 was used as a negative control because it is a transcription factor whose consensus binding element was also found commonly in the promoter regions of the neuroendocrine markers. **D.** LNCaP was treated with 10 μM enzalutamide and cells were harvested at the indicated time. Kinetics of NED markers and TCF4 transcription following enzalutamide treatment was analyzed using QPCR. The results demonstrated increased mRNA levels of TCF4 in 3 hours while the expression levels of NED markers (ChgA, NSE, and PTHrP) was induced at 24 hours. This observation suggests that TCF4 signals upstream of NED markers.

### Blocking TCF4/PTHrP partially reverses enzalutamide resistance in human CaP cell lines

To examine the biological significance of TCF4 and NED induction on enzalutamide resistance, we generated two additional human CaP cell lines that are resistant to enzalutamide (22Rv1-EnzR and VCaP-EnzR). Then, of the NED markers, we focused on PTHrP as a potential mediator of enzalutamide resistance because this peptide has been demonstrated to mediate castration resistance (18, 19). Consistent with the published data reporting that TCF4 is a binding partner of β-catenin (20, 21), pretreatment with the β-catenin degradation activator, XAV939 (22) at 10 μM five minutes prior to enzalutamide treatment, abrogated PTHrP mRNA induction by enzalutamide in all three human CaP cell lines (Fig 3A). In addition to the ligand, we assessed whether enzalutamide altered the expression of the PTHrP receptor, PTH1R. The result, shown in Fig 3B, revealed that 10 μM enzalutamide significantly increased mRNA levels in LNCaP, VCaP, and 22Rv1. Next, TCF4 and PTHrP were blocked in all three enzalutamide-resistant cell lines using the reported inhibitors – PKF118-310 and PTHrP_(7-34)_ (23, 24). Both PKF118-310 and PTHrP_(7-34)_ significantly decreased the cellular proliferation of all three cell lines in a concentration-dependent manner over a three-day period. Between the two inhibitors, PTHrP_(7-34)_ had a more moderate effect. In the three enzalutamide-resistant cell lines (LNCaP-EnzR, 22Rv1-EnzR, and VCaP-EnzR), PKF118-310 again inhibited the cellular proliferation in a concentration-dependent manner up to 50 μM after 3 days (Fig 3C). Similarly, PTHrP_(7-34)_ treatment also decreased cell count of LNCaP-EnzR, 22Rv1-EnzR, and VCaP-EnzR in a concentration-dependent manner up to 1 mM after 3 days. To assess whether PFK118-310 and PTHrP_(7-34)_ affected enzalutamide sensitivity, we next cultured the three enzalutamide-resistant cell lines with a fixed concentration of PKF118-310 (5 μM) or PTHrP_(7-34)_ (10 μM) and varying concentrations of enzalutamide (0-10 μM). The results demonstrated that both PKF118-310 and PTHrP_(7-34)_ reversed enzalutamide resistance in LNCaP-EnzR, 22Rv1-EnzR, and VCaP-EnzR (Fig 3D). It should be noted that enzalutamide still exhibited a concentration-dependent inhibitory effect on all three enzalutamide-resistant cell lines, demonstrating that these cells have a relative and not an absolute resistance to enzalutamide.

**Figure 3.**
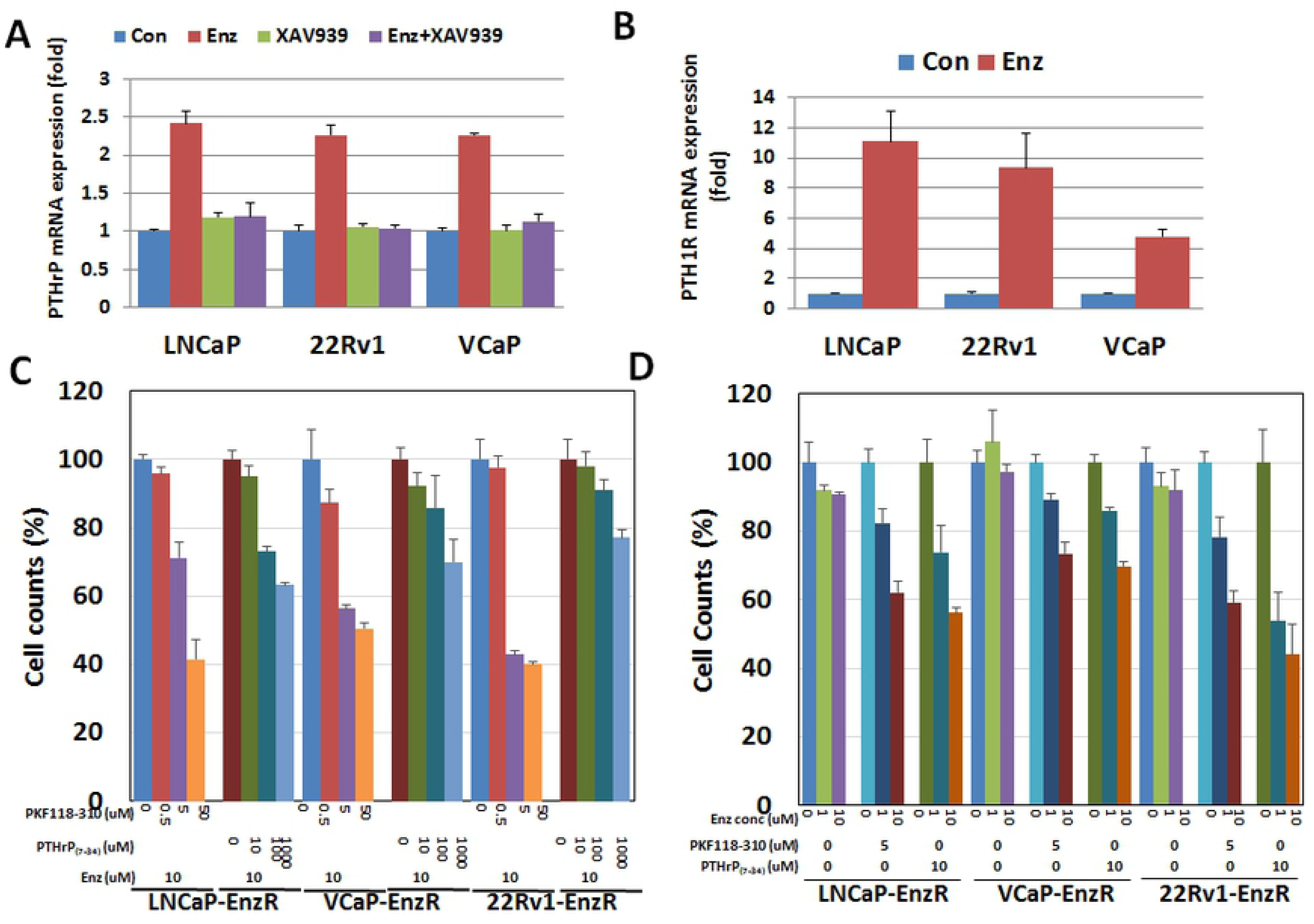
Effect of blocking TCF4/β-catenin (PKF118-310) or PTHrP (PTHrP_(7-34)_) on enzalutamide resistant prostate cancer cells. **A.** LNCaP, 22Rv1, and VCaP were treated with enzalutamide (10 μM) and/or XAV939 (10 μM), β-catenin degradation activator for 48 hours. XAV939 treatment was carried out 5 min prior to the addition of enzalutamide. PTHrP mRNA induction after 48 hours of treatment with 10 μM enzalutamide was completely blocked by XAV939, β-catenin inhibitor in LNCaP, 22Rv1 and VCaP. **B.** LNCaP, 22Rv1, and VCaP were treated with 10 uM enzalutamide for 48 hours. The PTHrP receptor, PTH1R, mRNA level significantly increased after enzalutamide treatment in all three cell lines. **C.** Three enzalutamide-resistant human prostate cancer cell lines (LNCaP-EnzR, 22Rv1-EnzR, and VCaP-EnzR) were treated with 10 μM enzalutamide and increasing concentrations of PKF118-310 and PTHrP_(7-34)_ as indicated. After 48 hours, viable cells were counted. In the presence of 10 μM Enz, PKF118-310 or PTHrP_(7-34)_ increased enzalutamide sensitivity in the enzalutamide resistant prostate cancer cell lines, LNCaP-EnzR, VCaP-EnzR, and 22Rv1-EnzR. **D.** Enzalutamide-resistant human prostate cancer cell lines were treated with a fixed concentration of PKF118-310 (5 μM) or PTHrP_(7-34)_ (10 μM) and varying concentrations of enzalutamide (0-10 μM). After 48 hours, viable cells were counted. In the presence of 5 μM of PKF118-310 or 10 μM of PTHrP_(7-34)_, enzalutamide inhibited cellular proliferation in a concentration-dependent manner.

Because TCF4 has been linked to Wnt signaling, we next examined the effect of typical canonical and non-canonical Wnts [Wnt1, 3b, 5a, and 5b (10 ng/ml)] on NED. The results demonstrated no significant effect of on the expression levels of ChgA, NSE, and PTHrP mRNA after 48 hours (Supp Fig 2). These results suggest that Wnts likely do not regulate NED in our experimental conditions and that the effect of TCF4 on NED is likely independent of the Wnt signaling pathway.

### In human CaP tissues, PTHrP and TCF4 co-localize and are associated with metastasis

To clinically validate these observations, we next carried out immunofluorescence (IF) microscopy on the CRPC tissue microarray (TMA) obtained from the University of Michigan’s rapid autopsy program. Although this CRPC TMA was established prior to the formal approval of enzalutamide by the United States Food and Drug Administration, co-localization of PTHrP and TCF4 was frequently observed in CRPC tissues (Fig 4A). PTHrP and TCF4 expression had positive correlation (Fig 4B). Furthermore, metastatic CaP was found to express higher levels of TCF4 and PTHrP (Fig 4C and D, respectively) when compared to localized CaP and benign prostate hyperplasia. Human kidney tissues were used as a positive control. As negative controls, tissues were stained only with the secondary antibody conjugated with FITC or red fluorescence protein (RFP).

**Figure 4.**
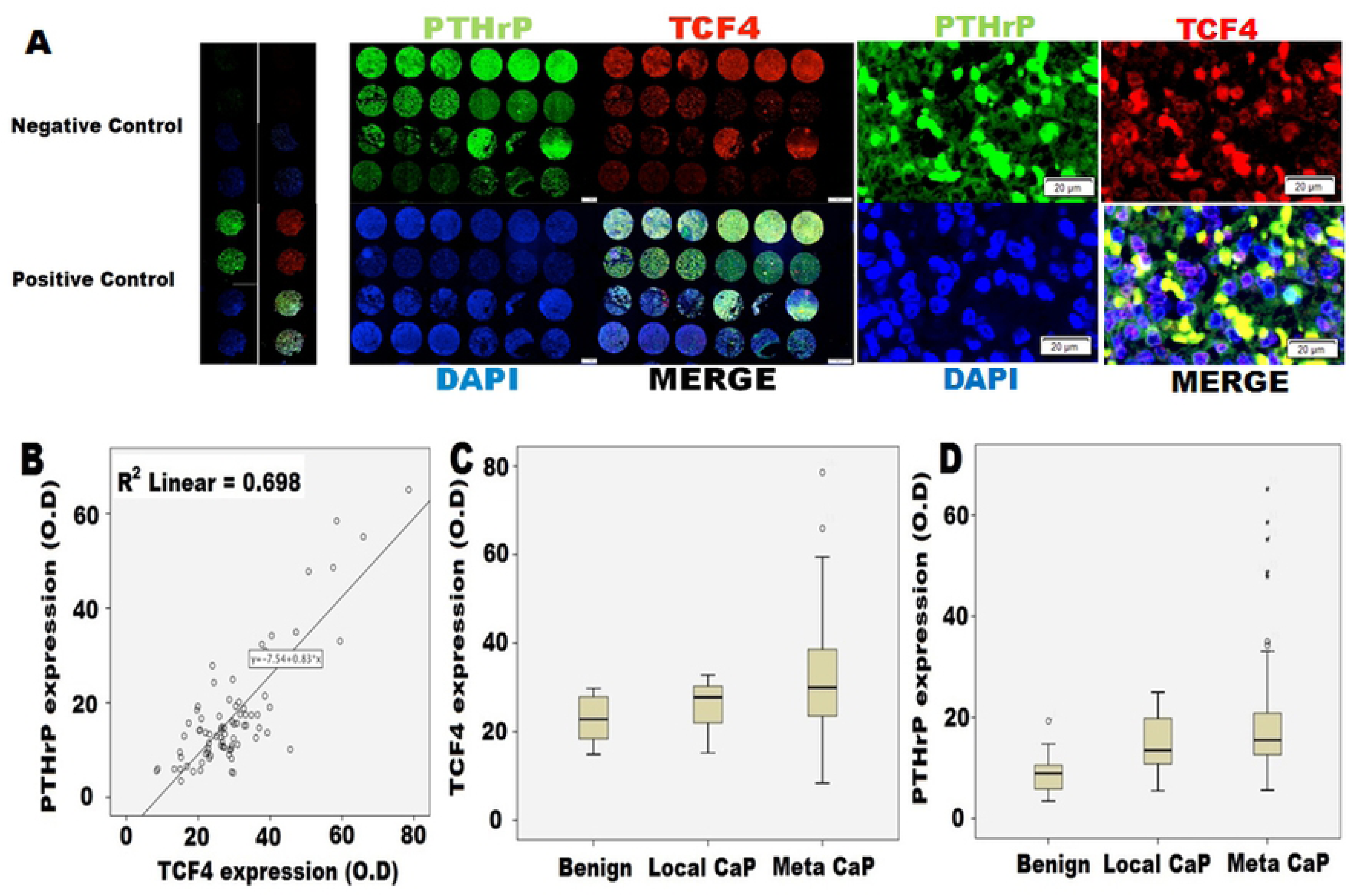
Human CRPC tissue microarray (TMA) analysis. TMA was obtained from the rapid autopsy program at the University of Michigan. This array contains 51 CRPC samples as well as 16 benign prostate tissues and 12 localized prostate cancer tissues for controls. **A.** Immunofluorescence microscopy demonstrated a consistent co-localization of TCF4 (red) and PTHrP (green) in CRPC tissues. **B.** There was a correlation between PTHrP and TCF4 expression. **C.** Protein expression levels of TCF4 and **D.** PTHrP in patients with localized CaP (Local CaP) and metastatic CaP (Meta CaP). TCF4 and PTHrP protein levels were higher in CaP when compared with benign and increased even further in metastatic group when compared with localized CaP group. Benign n=16, Local CaP n=12, meta CaP n=51.

### TCF4 has an oncogenic function

To assess the effect of TCF4 *in vivo*, we established a doxycycline-inducible TCF4-expressing cell line, LNCaP-TCF4. After screening multiple clones, the one with the highest induction level of TCF4 on treatment with 1 μg/ml of tetracycline was selected and further characterized (clone #3, Supp Fig 3A). TCF4 induction slightly increased cellular proliferation but the difference was not statistically significant (Supp 3B). As predicted based on our results, TCF4 induction resulted in a relative resistance to enzalutamide up to 10 μM when cultured over three days (Supp Fig 3C). In addition, tetracycline treatment stimulated the expression of NED markers, ChgA, NSE, and PTHrP (Supp Fig 3D).

When LNCaP-TCF4 was injected into flanks of Rag2-/-, γc-/-immunodeficient mice and TCF4 expression was induced by doxycycline following surgical castration, a relative resistance to enzalutamide treatment was observed over a six-week period (Fig 5A). There was no change in histology among the harvested tumors regardless of the grouping (Fig 5B). Immunofluorescence microscopy demonstrated that enzalutamide treatment or TCF4 induction (doxy) increased the expression of the NED marker, PTHrP (Fig 5C). In addition, co-localization of TCF4 and PTHrP was confirmed. Subsequently, immunoblot confirmed the immunofluorescence microscopy results in that increased TCF4 and PTHrP proteins were observed in groups treated with either enzalutamide or doxycycline (Fig 5D). QPCR demonstrated an increase in TCF4 and PTHrP mRNA levels upon treatment with enzalutamide as well as doxycycline (Fig 5E). Collectively, these results suggest that TCF4 renders CaP cells more aggressive.

**Figure 5.**
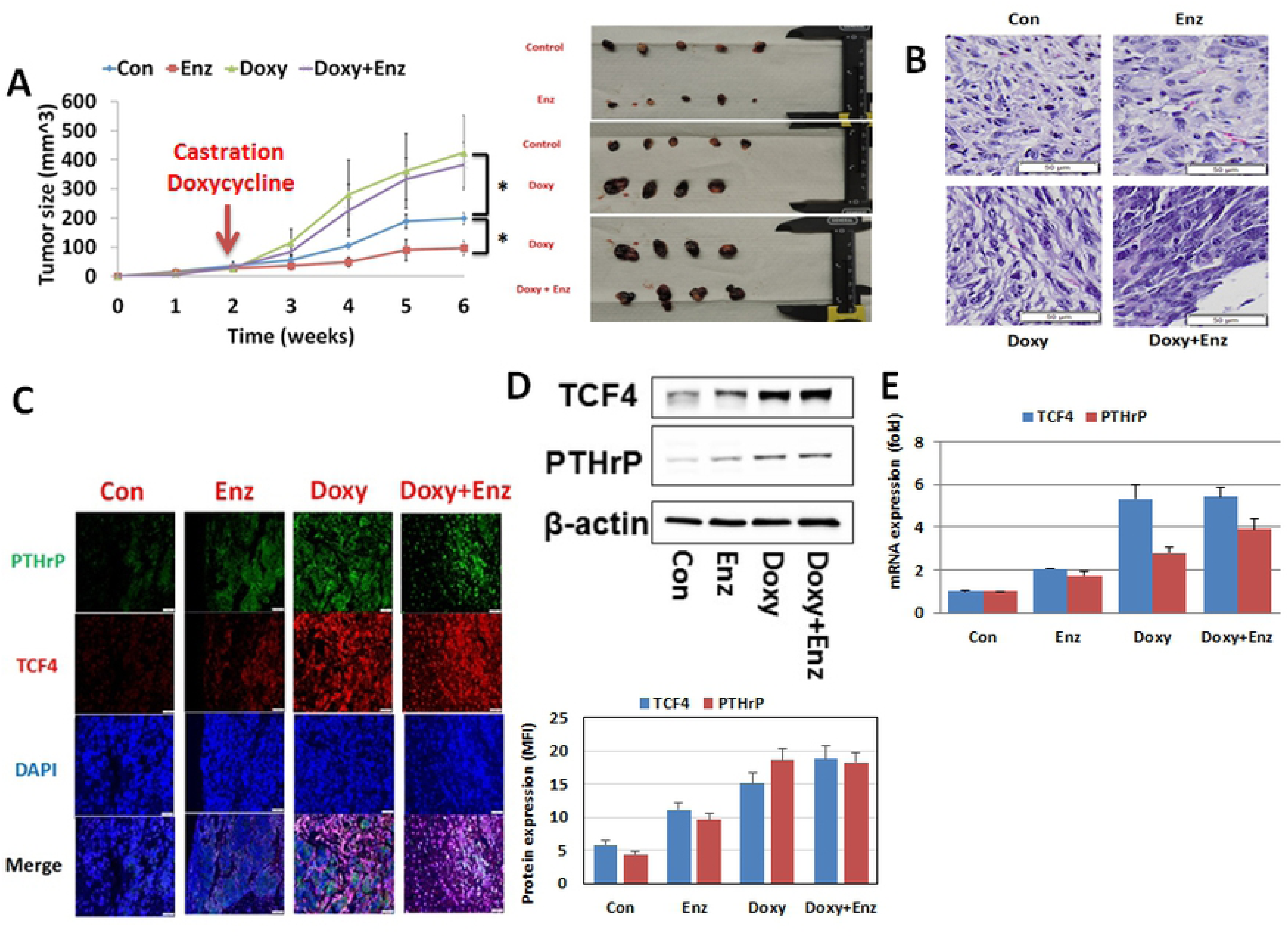
TCF4 induces enzalutamide resistance in the human prostate cancer cell line, LNCaP. LNCaP transfected with tetracycline-inducible TCF4 plasmid (LNCaP-TCF4) was injected into the flanks of twenty Rag2-/-, γ_c_-/- immunodeficient mice. When tumors reached an average size of 3 mm, the mice were divided into four groups of five each. Where indicated, 10 mg/kg enzalutamide was delivered orally daily. In the designated groups, TCF4 was delivering doxycycline via the drinking water. At the end of the indicated duration, all tumors were harvested and analyzed for protein and mRNA expression. **A.** When TCF4 expression was induced with doxycycline (Doxy), tumor growth rate increased when compared to the control group. In the absence of TCF4 induction, enzalutamide treatment slowed tumor growth rate. However, enzalutamide treatment had no demonstrable inhibitory effect in TCF4-induced doxycycline group. Con = LNCaP-TCF4 without doxycycline. Enz = enzalutamide. **B.** H&E staining. There was no difference among all groups. **C.** Immunofluorescence staining for PTHrP (green), TCF4 (red) with DAPI (blue) staining. Increased TCF4 and PTHrP protein levels were observed following the induction of TCF4 with doxycycline. Directly supporting the tissue culture data, enzalutamide treatment also increased protein levels of TCF4 and PTHrP. **D and E**. Effect of enzalutamide and TCF4 overexpression on TCF4 and PTHrP by western blot analysis (**D)** and QPCR (**E**). Immunoblot and QPCR both demonstrated increased expression of TCF4 and PTHrP following TCF4 induction (Doxy). A more modest increase in TCF4 and PTHrP expression was observed with enzalutamide (enz) treatment. Error bars indicate average ± SE and * p-value<0.05.

### TCF4 inhibitor and PTHrP antagonist inhibit the proliferation of enzalutamide resistant prostate cancer cells *in vivo*

To study the therapeutic potential of targeting the TCF4/PTHrP axis, we carried out an *in vivo* study with LNCaP-EnzR in mice. After establishing tumor xenografts, all animals underwent a bilateral orchiectomy and were administered 10 mg/kg of enzalutamide via oral gavage daily. To predesignated groups, 0.85 mg/kg of PKF-118-310, 0.2 mg/kg of PTHrP_(7-34)_, or both in combination were delivered daily. At the end of seven weeks, the results demonstrated that PKF118-310 and PTHrP_(7-34)_ dramatically when combined with enzalutamide decreased the tumor xenograft growth dramatically compared to that of enzalutamide monotherapy (control) over a seven-week period (Fig 6A). However, there were no synergistic effect between PKF118-310 and PTHrP antagonist, PTHrP_(7-34)_. Again, H&E staining showed no significant changes in the histology of the treated xenografts (Fig 6B). Immunofluorescence microscopy confirmed the co-localization of TCF4 and PTHrP while MFI measurement demonstrated that PKF118-310 treatment decreased PTHrP protein levels (Fig 6C). However, PTHrP_(7-34)_ treatment again had no effect on TCF4 expression. Also consistent with the mechanism of action of PKF118-310 in which the interaction between TCF4 and B-catenin is disrupted, PKF118-310 treatment did not alter the expression levels of TCF4. These observations collectively support our concept that TCF4 is an upstream signaling molecule of PTHrP. Supporting the immunofluorescence MFI result, immunoblot revealed a decrease in PTHrP protein levels following PKF118-310 but not PTHrP_(7-34)_ treatment (Fig 6D) levels. Again, no obvious changes in TCF4 protein level was seen with either PFK118-310 or PTHrP_(7-34)_ administration. Result of the mRNA quantitation for TCF4 and PTHrP was consistent with that of immunoblot (Fig 6E).

**Figure 6.**
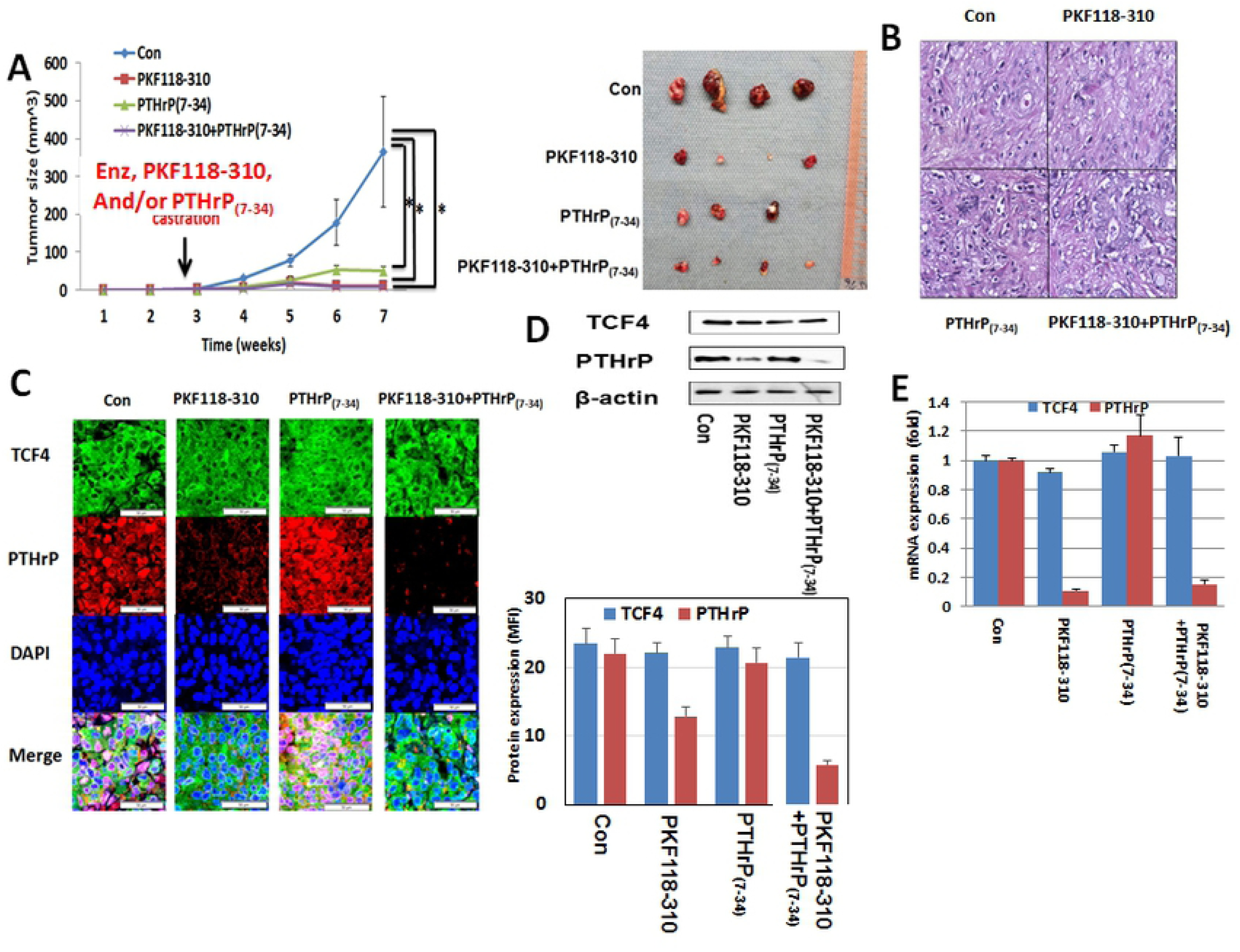
Effect of TCF4/β-catenin inhibitor (PKF118-310) and PTHrP antagonist (PTHrP_(7-34)_) in enzalutamide-resistant prostate cancer. After injection of LNCaP-EnzR into the flanks of forty Rag2-/-, γ_c_-/- immunodeficient mice, all mice were surgically castrated divided into four groups of ten each. Animals in predesignated groups were treated daily with PKF118-310 (0.85 mg/kg intraperitoneal) and/or PTHrP_(7-34)_ (0.2 mg/kg subcutaneous). All mice were administered daily 10 mg/kg enzalutamide orally. **A.** Treatment of PKF118-310 and/or PTHrP_(7-34)_ with 10 mg/kg enzalutamide decreased tumor growth compare with vehicle treatment control group (con). **B.** H&E staining. There was no difference among all groups. **C.** Immunofluorescence staining for TCF4 (green), PTHrP (red) with DAPI (blue) staining. Consistent with its mechanism of action, PFK118-310 treatment decreased PTHrP protein levels. However, there was no effect on TCF4 levels. In contrast, PTHrP_(7-34)_ had no demonstrable effect on the protein levels of both TCF4 and PTHrP. Treatment of PKF118-310 decreased PTHrP protein (**D**) and mRNA expression (**E**). Error bars indicate average ± SE and * p-value<0.05.

Finally, PKF118-310 and PTHrP_(7-34)_ as a monotherapy (without enzalutamide) at the same dosage had a more moderate effect on the growth of LNCaP-EnzR xenografts over a six-week period (Supp Fig 4A). Compared to the vehicle only control group, enzalutamide treatment slightly decreased the tumor growth rate. But this difference was not statistically significant. Again, H&E staining demonstrated no obvious change in histology (Supp Fig 4B). As seen with above results, immunofluorescence microscopy and MFI showed that PKF118-310 decreased PTHrP but not TCF4 protein levels (Supp Fig 4C). These observations were supported by the immunoblot (Supp Fig 4D) and QPCR (Supp Fig 4E). These results collectively suggest that TCF4 and PTHrP inhibitor may be an effective treatment option in the enzalutamide resistant CaP cells.

## DISCUSSION

In the present study, we investigated the mechanism of enzalutamide resistance in CaP. After establishing multiple cell lines that are resistant to enzalutamide, NED was identified as a significant event through RNAseq. Subsequently, the transcription factor TCF4 was found to regulate NED which in turn, rendered the cells resistant to enzalutamide in part via PTHrP. *In vivo* studies confirmed the critical role of TCF4 in enzalutamide resistance and NED. Collectively, these observations demonstrate that enzalutamide stimulates TCF4 transcription which then leads to NED and treatment resistance. More importantly, the *in vivo* studies suggest that TCF4 may potentially be a new therapeutic target in patients with CRPC.

Castration resistance is the ultimate clinical feature of lethal CaP. Despite being resistant to castration, CRPC usually has a clinically significant androgen signaling pathway. This apparent paradox is due to alterations in intracellular androgen synthesis that permit CaP cells to survive and proliferate under low extracellular androgen concentrations (25-27). Indeed, the presence of such mechanism in CRPC has been validated clinically with agents such as abiraterone acetate and enzalutamide that target androgen biosynthesis and androgen receptor, respectively (28-30).

In treating men with CRPC, enzalutamide is a key second generation anti-androgen used widely due to its ease of administration as well as a low toxicity profile (6). Notwithstanding, as with all second-generation androgen manipulations, resistance to enzalutamide treatment emerges inevitably with a median time of 4.8 months (31). Although the precise sequence of molecular alterations that result in enzalutamide resistance in CaP remains unclear, it is likely that the mechanism is heterogeneous and involves perturbation of multiple signaling pathways and factors (32-40). One major proposed mechanism is the aberrant expression of AR splice variants, especially AR-V7 (32-34). It has been reported that the level of AR-V7 expression positively correlates with enzalutamide resistance in CaP (32). Another commonly proposed mechanism of enzalutamide resistance is neuroendocrine trans-differentiation. (37, 38, 41). Specifically, enzalutamide treatment has been associated with the dysregulated transcription factors and genetic abnormalities that lead to NED (42-44). A third class of mechanism of enzalutamide resistance is AR point mutation. Among several well-characterized AR point mutations, T878A (previously T877A) and F877L (previously F876L) have been reported to increase enzalutamide resistance in CaP cell lines (35, 36). However, AR point mutations do not appear to be a major mechanism of enzalutamide resistance in our experimental models as AR mRNA was completely sequenced in all three enzalutamide-resistant cell lines. Rather, our results suggest NED as the relevant mechanism of enzalutamide resistance in CaP and propose TCF4 as a key molecule in mediating NED in enzalutamide-resistant CaP cells.

Although TCF4 is a transcription factor that has been implicated in the canonical β-catenin-mediated Wnt signaling (10), present results suggest a new function that is independent of Wnt signaling: regulation of NED. This activity of TCF4 is dependent on the interaction with β-catenin as confirmed by the results of the β-catenin degradation activator study (XAV939) (Fig. 3A). However, treating CaP cells with both classical and non-classical Wnts did not induce NED. In addition, it should be noted that NED in the context of CaP is likely different than the class small cell neuroendocrine prostate cancer (NEPC) cells. For example, although NEPC do not express AR (45), cells with NE phenotype in CaP have been reported to express AR (46, 47). This observation is consistent with the concept of transdifferentiation in which in CaP, adenocarcinoma cells differentiate into cells expressing NE markers (48, 49). Indeed, Lin and colleagues have proposed the concept of NE-like CaP cells that have retained some of the features of CaP adenocarcinoma cells while attaining NE features (50). Regardless, it has recently been reported that NED accounts for approximately 25% of lethal prostate cancer (51). Whether TCF4 is involved in all NE-like CaP requires further investigation.

The current study also shed additional light on the mechanistic link between NED and enzalutamide resistance. Using RNAseq as well as overexpression and neutralization studies, we have observed that the transcription factor TCF4 mediates NED upon enzalutamide treatment. Of the NED markers, we have focused on PTHrP for the subsequently functional studies because PTHrP has previously been reported to mediate castration resistance (19). In this regard, our data also support this concept as neutralization of PTHrP reversed enzalutamide resistance. In addition to blocking PTHrP, we have observed that the TCF4 inhibitor (PKF118-310) increased enzalutamide sensitivity and decreased PTHrP protein levels in enzalutamide resistant CaP cell lines in tissue culture and *in vivo*. Therefore, disrupting the NED axis by targeting either TCF4 or PTHrP may be an effective therapeutic approach in treating CRPC.

Despite the potentially significant implications of the current study, our findings should be accepted with caution for the following reasons. First, there are multiple second-generation anti-androgens. Thus, additional studies are needed to determine whether TCF4-mediated NED is a class-specific effect. Second, the molecular connection between TCF4 and enzalutamide is not clear. Currently, we are exploring a potential interaction between androgen receptor and TCF4.

In conclusion, the present study suggests that TCF4 mediates enzalutamide resistance via a Wnt-independent pathway. In addition, we have observed that blocking NED via TCF4 or PTHrP inhibitors is potentially therapeutic. In the future, we plan to continue uncovering the molecular link between enzalutamide and TCF4/NED as well as further characterizing the therapeutic potential of blocking TCF4 or PTHrP.

